# A Reinforcement Learning Framework for Pooled Oligonucleotide Design

**DOI:** 10.1101/2021.08.25.455853

**Authors:** Benjamin M. David, Ryan M. Wyllie, Ramdane Harouaka, Paul A. Jensen

**Affiliations:** Department of Bioengineering, University of Illinois at Urbana-Champaign; Biotechnology and Bioengineering Department, Sandia National Laboratories, Livermore, CA; Department of Microbiology, University of Illinois at Urbana-Champaign; Carl R. Woese Institute for Genomic Biology, University of Illinois at Urbana-Champaign

## Abstract

The goal of oligonucleotide (oligo) design is to select oligos that optimize a set of design criteria. Oligo design problems are combinatorial in nature and require computationally intensive models to evaluate design criteria. Even relatively small problems can be intractable for brute-force approaches that test every possible combination of oligos, so heuristic approaches must be used to find near-optimal solutions. We present a general reinforcement learning framework, called OligoRL, to solve oligo design problems with complex constraints. OligoRL allows “black-box” design criteria and can be adapted to solve many oligo design problems. We highlight the flexibility of OligoRL by building tools to solve three distinct design problems: 1.) finding pools of random DNA barcodes that lack restriction enzyme recognition sequences (CutFreeRL); 2.) compressing large, non-degenerate oligo pools into smaller degenerate ones (OligoCompressor); and 3.) finding Not-So-Random hexamer primer pools that avoid rRNA and other unwanted transcripts during RNA-seq library preparation (NSR-RL). OligoRL demonstrates how reinforcement learning offers a general solution for complex oligo design problems. OligoRL and its associated software tools are available as a Julia package at http://jensenlab.net/tools.

## 1 Introduction

Synthesized oligonucleotides (oligos) are a ubiquitous tool in molecular biology. Researchers use multiple design criteria for oligos or oligo pools based on the problem they are trying to solve. Such criteria include alignment to a DNA sequence or genome, avoidance of sequences or genomes, GC content, melting temperature, or degeneracy (the number of unique sequences in an oligo pool defined by degenerate base codes (Figure 1A))[1]–[3]. Many oligo pools are designed by a process of elimination. All possible oligos are considered at the start, and the pool is narrowed down until only the candidates that satisfy all criteria remain[4]–[6]. This strategy may be effective for small problems, but it does not scale well. Increasing the length of an oligo or adding degenerate bases exponentially increases the size of an oligo pool. For example, using only the non-degenerate bases `A`, `C`, `G`, or `T`, there are 4,096 unique DNA hexamer (6-mer) sequences but 16,777,216 unique 12-mer sequences. If all 15 degenerate bases are allowed, there are 11,390,625 hexamer and 129,746,337,890,625 12-mer sequences[7]. Interactions among short sequences can also lead to combinatorial complexity. For example, designing pools of primers for multiplex PCR amplification requires avoiding primers that form heterodimers. Adding any oligo requires testing for sequence overlap with all other oligos, creating dependencies between all candidates.

**Figure 1:**
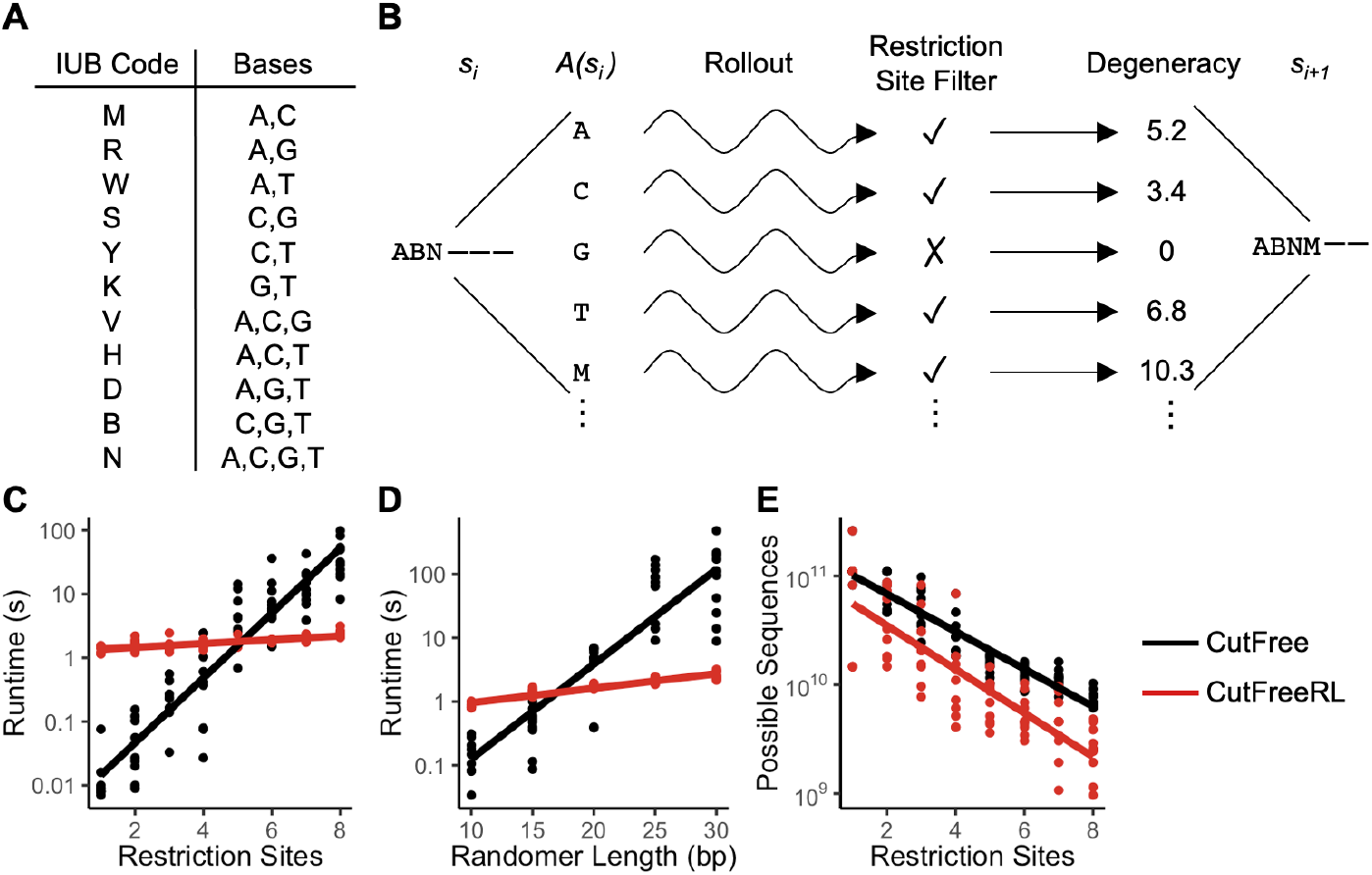
CutFreeRL was developed using the OligoRL framework. **A**. IUB codes specify all combinations of the bases `A`, `C`, `G`, and `T`. **B**. An OligoRL agent in state *s*_*i*_ selects an action *a*_*i*_ from the set of available actions *A*(*s*_*i*_) in order to move to state *s*_*i*+1_. A reward *r*_*i*_ is associated with the selected action. Rollout uses simulations to find the action with the highest expected total reward. The process repeats from state *s*_*i*+1_ onward until the entire oligo has been selected. CutFreeRL’s reward function implements a restriction site filter. Oligos that do not contain an unwanted restriction site are scored in proportion to their degeneracy. **C**. CutFreeRL exhibits better scaling than the original, MILP-based CutFree algorithm as the number of blocked restriction sites increases. **D**. CutFreeRL solve problems with longer randomers faster than CutFree. CutFreeRL also has less variability in runtimes for large problems. **E**. Randomers designed with CutFreeRL have lower degeneracy than those designed with CutFree, although the rate of decrease in degeneracy as restriction sites are blocked is the same for both algorithms.

A process of elimination creates oligo pools that pass a set of design filters; however, these pools are rarely optimal with respect to the design criteria. There are two obstacles to designing optimal pools. First, the combinatorial structure of oligo design problems makes these problems computationally intractable. Heuristics or sophisticated approximation strategies must be used to find near-optimal solutions since brute-force techniques are impractical. Second, incorporating a traditional optimization framework requires that the design criteria be expressed in a specific way, e.g. as linear, convex, or mixed-integer constraints. Many oligo design criteria cannot be written in this way. For example, the melting temperature of an oligo is a nonlinear, discontinuous function of oligo structure[8], and counting the number of times an oligo binds to a genome would require millions or billions of integer constraints depending on the size of the genome.

Here we describe a general reinforcement learning framework—OligoRL— for designing oligonucleotide pools. OligoRL finds near-optimal (within 10% of the optimal objective value) oligo pools using “black-box” design criteria. We demonstrate the generality of OligoRL by solving three distinct oligo design problems: 1.) finding degenerate barcodes that lack restriction enzyme recognition sites; 2.) compressing a set of individual oligos into a smaller pool of degenerate oligos; and 3.) creating semi-random hexamer libraries that avoid rRNA when preparing RNA-seq libraries. OligoRL solves all three problems using the same rollout algorithm [9], [10]. (Only the reward functions and action spaces are changed to match the design criteria for each problem.) Thus, OligoRL can be used to solve many challenging oligo design problems, including problems with design criteria that are difficult to represent algebraically.

## 2 Results

### 2.1 The OligoRL Framework

OligoRL formulates the oligo design problem as a Markov Decision Process (MDP)[11]. The MDP describes how an agent in a state *s*_*i*_ selects an action *a*_*i*_ that moves the agent to a new state *s*_*i*+1_ (Figure 1B). The transition between states is accompanied by a reward *r*_*i*_. The agent’s goal is to select actions that maximize the sum of all the rewards. Our problem is to build an oligo of length *L* by selecting degenerate base codes at each position. The oligo codes are selected sequentially beginning at the 5’ end. An agent in state *s*_*i*_ has selected the first *i* − 1 oligo codes, so the agent begins at state *s*_1_, when zero oligo codes have been selected, and finishes at state *s*_*L*+1_. The state defines not only how many but also which codes have been selected. An agent that has selected codes `ACG` is in a different state than an agent that has selected codes `ACT`.

Once in state *s*_*i*_, the agent selects the code to place at position *i*. This selection corresponds to the action *a*_*i*_, which is drawn from the set of possible codes *A*(*s*_*i*_). The set of allowed codes is state-dependent—the codes selected for the prior positions 1 … *i* − 1 can change the codes available to the agent at position *i*. Each available code *a*_*i*_ ∈ *A*(*s*_*i*_) has an associated reward *r*_*i*_(*a*_*i*_). This reward depends on the entire oligo up to and including position *i*. The final reward *r*_*L*_(*a*_*L*_) is based on the entire oligo.

We do not make any assumptions about the reward functions. For example, the reward for an oligo can be based on aligning the oligo to a genome and counting the number or quality of the hits. It is also possible to set all but the final rewards to zero, delaying the reward calculation until the entire oligo has been selected. Furthermore, the reward function can be applied to either a single oligo or an entire oligo pool. The flexibility of the reward function underlies the generality of our approach, but it also requires us to solve the oligo selection problem by simulation.

We use a rollout algorithm to choose the best code at each position. Rollout is a reinforcement learning (RL) technique used to solve large MDPs by simulating trajectories using a computer model [9], [10]. In state *s*_*i*_ we begin by considering the first code *a*_1_ ∈ *A*(*s*_*i*_). We simulate ahead to the end of the oligo, choosing codes randomly and summing the rewards. By averaging the rewards from many random trajectories, all beginning with action *a*_1_, we can estimate the average reward the agent will experience when code *a*_1_ is selected. We compute this reward-to-go estimate for all other actions available at state *s*_*i*_. The action we ultimately choose in state *s*_*i*_ corresponds to the maximum reward from the rollout simulations. After the code is selected, we move to the next position (state *s*_*i*+1_) and repeat the rollout process starting at the new state.

To demonstrate the flexibility of OligoRL, we apply the framework to three pooled oligo design problems. First, we design pools of random DNA barcodes that lack restriction enzyme recognition sequences. We previously solved this problem using a mathematical programming algorithm called CutFree[3]. The new OligoRL implementation returns near-optimal solutions but is more computationally efficient than CutFree on large problems.

Next, we use OligoRL to implement an oligo compressor. Given a large pool of non-degenerate oligos, the compressor finds a smaller pool of degenerate oligos that contains every sequence (and only those sequences) in the larger pool.

Finally, we use OligoRL to design pools of Not-So-Random (NSR) primers. NSR pools are used to selectively prime reverse transcription in when preparing sequencing libraries. The original NSR pools were designed to avoid hybridization to rRNA transcripts[4]. Using OligoRL, we found smaller NSR pools with increased uniformity across all mRNAs in a representative organism. This final example demonstrates “black-box” reward functions that map the NSR primers to transcriptomes and calculate the uniformity of an NSR pool. Neither of these reward functions can be expressed as algebraic constraints on the OligoRL problem.

### 2.2 CutFreeRL

DNA barcodes are pools of randomers used to label individual genetic parts. Random barcodes can be problematic for restriction enzyme-based cloning methods since some of the random sequences will contain restriction enzyme recognition sites. The unwanted sites in the barcodes can lead to incorrect DNA constructs or incorrect barcoding.

We previously developed the CutFree algorithm to design degenerate oligo pools that are free from a user-specified set of restriction sites[3]. CutFree is a mixed integer linear program (MILP) that uses constraints to guarantee that the restriction sites do not appear in the resulting degenerate randomers. Users input a set of unwanted restriction sites and the degenerate codes allowed at each position. CutFree uses an MILP solver to maximize the degeneracy of the randomer subject to the restriction site constraints. Unfortunately, the runtime of CutFree increases exponentially when increasing either the number of blocked restriction sites or the length of the randomer. CutFree’s MILP framework also causes large variations in runtime, as the computational difficulty of an MILP can change drastically when even small changes are made to the objective or the constraints. Although CutFree can find provably optimal solutions for many small problems, the algorithm is intractable for large problems with long randomers or many blocked restriction sites.

We used our reinforcement learning framework to develop CutFreeRL, a rollout-based alternative to the CutFree algorithm. Like CutFree, CutFreeRL creates a single degenerate oligo. Beginning at the 5’ end, the agent selects a degenerate base *a*_*i*_ in state *s*_*i*_ to maximize the expected future reward. As we showed with CutFree, maximizing diversity of the randomer is equivalent to maximizing the sum of the log-degeneracy of each individual base[3], so *r*_*i*_ = log *n*_*b*_ where *n*_*b*_ is the degeneracy of base *b*. (For example, *n*_A_ = *n*_G_ = *n*_C_ = *n*_T_ = 1, *n*_M_ = 2, and *n*_N_ = 4.) Before selecting a base, the agent checks that no unwanted restriction site appears in positions 1 … *i* of the oligo. Unlike the MILP-based CutFree, the check for restriction sites is not encoded algebraically. Instead, CutFreeRL simply calls a function that matches the unwanted sites against the oligo. The reward for a randomer is set to zero if a simulated oligo contains to any of the restriction sites.

We benchmarked CutFreeRL against the original CutFree algorithm by designing randomers of different lengths or numbers of restriction sites. Restriction sites were selected randomly from the set of all commercially available, palindromic, six base pair restriction enzyme recognition sequences in REBASE[12]. As previously shown, CutFree’s runtime increased exponentially when either the randomer length or the number of restrction sites increased (Figure 1C,D).

Conversely, the runtime for CutFreeRL varied linearly with respect to randomer length and the number of restriction sites. This linear response was expected since the rollout algorithm uses a fixed horizon to check if a randomer contains any restriction sites. The horizon extends from the current base back as far as the longest restriction site. Like most MILPs, the original CutFree algorithm scaled exponentially with either the randomer length or the number of restriction sites. For problems with more than five restriction sites and randomers with more than 17 bases, CutFreeRL was faster than the original CutFree algorithm.

Another disadvantage of the original CutFree MILP approach is that the runtimes can change unpredictably when different restriction sites are blocked from randomers of the same length. Rollout-based algorithms have more predictable runtimes than MILPs, and we observed less runtime variation between replicates for CutFreeRL than for the original CutFree approach.

While the original CutFree MILP framework frequently returns a provably optimal solution, CutFreeRL relies on random simulation to find a solution. CutFreeRL is limited to finding near-optimal solution since the rollout algorithm cannot visit the entire combinatoric search space. While the CutFreeRL approach sacrifices returning an optimal solution in order to handle solving larger problems, it returns randomers with degeneracy scores on the same order of magnitude as CutFree (Figure 1E). As expected, the degeneracies of oligos found by both approaches decreased exponentially as the number of restriction sites increased; however, there was no significant interaction between the number of restriction sites and the choice of method (CutFree or CutFreeRL), indicating that both CutFree and CutFreeRL scale similarly (*p* > 0.085, *t*-test).

### 2.3 OligoCompressor

Experimenters may have preexisting pools of oligos for a genomic assay. For example, Not-So-Random (NSR) reverse transcription primers selectively enrich mRNA transcripts when creating RNA-seq libraries[4]. Such pools bypass expensive and time consuming rRNA depletion steps. The cost of a set of oligos depends on the number of synthesis reactions. Synthesizing each oligo separately is often cost-prohibitive for large pools like NSR RNA-seq primers. However, synthesizing an oligo containing degenerate bases costs the same as synthesizing a single oligo sequence. It is possible that a set of oligos can be reduced to fewer synthesis reactions by compressing together similar sequences using ambiguous base codes. The resulting pool would be made up of degenerate oligos that encode the same information as the original uncompressed pool. Depending on the degree of similarity within sequence subsets, compressed oligo pools could substantially reduce synthesis costs.

We used the OligoRL framework to develop OligoCompressor. Given a pool of oligo sequences, OligoCompressor finds a smaller pool of degenerate oligos that contains the same sequences. By defining the reward function and action spaces, we ensure that OligoCompressor finds the smallest equivalent pool without introducing sequences not found in the original pool. The compressor follows a stepwise greedy approach where the agent is rewarded for maximizing the number of oligos from the original pool it can capture within a single degenerate oligo (Figure 2A). The agent is also constrained such that it can only capture oligos from the original pool, with prohibitively large penalties for incorporating sequences not found in the original pool. After a degenerate oligo has been selected, all of the sequences captured by that oligo are removed from the original pool, and the oligo selection process repeats until all sequences from the original pool have been captured.

**Figure 2:**
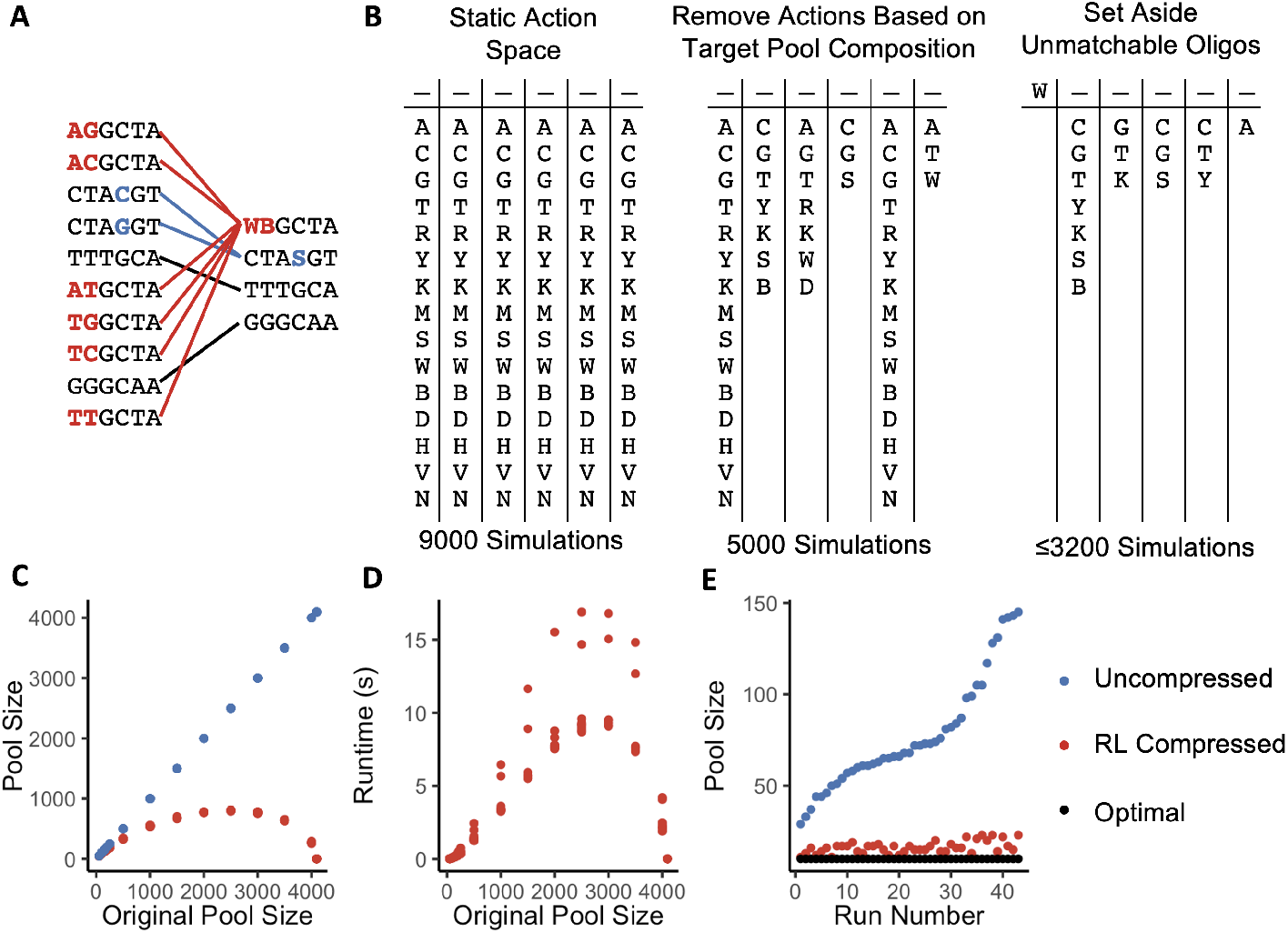
OligoCompressor reduces a large non-degenerate oligo pool into a smaller degenerate pool. **A**. OligoCompressor builds a degenerate oligo that captures the maximum number of oligos from the target pool. The selection process continues until all target oligos have been captured. **B**. OligoCompressor adjusts the action space after each base is selected to eliminate unnecessary simulation. After each base is selected, actions that do not correspond to partially-matched oligos are removed. **C**. OligoCompressor can compress pools of random hexamer sequences. A pool with all 4,096 possible hexamers is compressed down to a single degenerate oligo: `NNNNNN`. **D**. OligoCompressor builds one degenerate oligo at a time, so the algorithm’s runtime is proportional to the number of oligos in the final compressed pool. Runtimes are shown for ten random pools at each original pool size. **E**. OligoCompressor’s efficiency was tested by decompressing and re-compressing randomly generating degenerate pools of 10 oligos. The decompressed pools contained between 30–150 sequences.

OligoCompressor implements a dynamic action space to improve performance (Figure 2B). Since the sequences that OligoCompressor is trying to capture are known *a priori*, we can limit the action choices at each nucleotide position to only the bases in the target pool at that position. For example, if every remaining oligo in the target pool has either an `A` or `T` in the final position, then the agent doesn’t need to waste simulations by testing oligos that end in `G` or `C`. In this scenario, we can reduce the size of the action space for the agent at the final nucleotide position from 15 degenerate base codes down to three (`A`, `T`, or `W`). Furthermore, as the agent selects nucleotides, it commits to matching only a subset of the original pool. For example, an agent that selected a `W` for the first position can ignore any oligos in the target pool that do not start with an `A` or `T`. (The oligos that begin with `G` or `C` will need to be matched by a subsequent degenerate oligo.) Limiting the available actions and setting aside the unmatchable oligos quickly eliminates unnecessary simulations and reduces OligoCompressor’s runtime.

To assess the performance of OligoCompressor, we randomly generated hexamer pools of varying size and compressed them using the OligoRL framework. The degree of compression generally increased with the size of the starting pool (Figure 2C). This is expected since a pool of 1,000 random oligos is more likely to contain groups of similar, “compressible” oligos than a smaller pool of 100 oligos. We observed that OligoCompressor became increasingly effective when the number of the oligos in the pool approached the upper bound of 4,096 possible hexamer sequences. At approximately 2,500 starting oligos, the compressed pool size peaked around 800 oligos. Starting pools with more than 2,500 oligos could be compressed into even fewer oligos. As an extreme case, running OligoCompressor on a starting pool with all 4,096 possible hexamers returned the single degenerate oligo: `NNNNNN`.

OligoCompressor iterates by adding a single degenerate oligo until it captures every oligo in the starting pool. Its runtime therefore depends on both the number of iterations and the difficulty of finding a valid oligo during each iteration. The runtimes for OligoCompressor followed the same pattern as the degree of compression (Figure 2D). Pools with more than 2,500 starting oligos, for example, compressed faster because they produced fewer, more degenerate oligos and required fewer iterations to capture the starting pool. Interestingly, there was a quadratic relationship between runtime and pool size for small pool sizes (< 500 oligos). Since the degree of compression is relatively low for these smaller pools, the algorithm uses more iterations and must check itself against a larger fraction of the starting pool during each iteration. The relationship between runtime and pool size became linear for larger pools because each oligo selection iteration removed a larger fraction of the starting pool, reducing both the number of iterations and the runtime of each iteration.

We performed simulations to test how efficiently OligoCompressor can reduce pools of hexamers. We randomly selected degenerate, pre-compressed pools of 10 oligos of varying degeneracy and expanded the pools into a non-degenerate starting pool. The expanded pools contained between 30–150 oligos. We then used OligoCompressor to re-compress the expanded pools and compared the size of the re-compressed pools to the size of the pre-compressed pools (Figure 2E). On average, OligoCompressor returned a solution that contained 16.8 ± 3.7 oligos, while the largest solution contained 23 oligos for an expanded pool of 145 oligos. These results suggest that like CutFreeRL, OligoCompressor returns near-optimal solutions.

OligoCompressor excels at compressing pools of short oligos such as NSR primer pools used in single-cell RNA-seq. The compressed pools are equivalent to the uncompressed pools but cost less to manufacture. For example, a recently published NSR pool for mammalian single-cell RNA-seq contains 408 non-degenerate oligos[13]. OligoCompressor reduced the same set of sequences to 262 degenerate oligos—a savings of 36%. We can also use the OligoRL framework to directly design pools of NSR primers and tune the pool to specific design criteria.

### 2.4 NSR-RL

CutFreeRL and OligoCompressor use straightforward reward functions (sequence degeneracy and number of oligos matched) to search for optimal solutions. Both algorithms incorporate sequence alignment to find restriction sites or to compare short oligo sequences. As a final example, we use OligoRL to find oligo pools using a complex, multivariate reward function that uses a computationally-intensive sequence search across an entire transcriptome. Our goal is to find an optimal pool of Not-So-Random (NSR) primers that 1.) avoid rRNA, tRNA, or transcripts from any unwanted genes, 2.) bind to every gene in a target set at least once, 3.) uniformly cover the transcripts from targeted genes, and 4.) use the smallest number of oligos necessary to meet objectives 1–3.

Highly abundant transcripts from rRNA and tRNA genes constitute up to 95% of the RNA in the cell[14]. If these transcripts are not removed before sequencing, they can vastly inflate the sequencing cost needed to quantify the abundance of mRNA. In bulk RNA from eukaryotes, mRNA transcripts can be enriched by polyT selection; however, prokaryotic mRNAs are not polyadenylated, and the highly abundant rRNA transcripts must instead be removed by physical capture with silica columns or magnetic beads[15]–[17]. Column- and bead-based separations are not possible in single-cell RNA-seq studies where libraries are prepared from picograms of RNA in droplets or microwells.

Some RNA-seq protocols use random hexamers to prime reverse transcription. The hexamers bind randomly across the transcriptome, so libraries made from total RNA will be dominated by rRNA. Recently, NSR primers have been used to selectively amplify non-rRNA sequences[4]. An NSR pool contains only the hexamers that are not found in the rRNA genes, so rRNA transcripts are not primed for reverse transcription and subsequent amplification.

Current workflows for designing NSR hexamer primers start with a pool of all 4,906 possible hexamers and remove hexamers that appear in the undesired transcripts. The remaining hexamers are aligned to the rest of the transcriptome. The OligoRL framework can improve NSR selection in two ways. First, OligoCompressor can reduce the cost of a brute-force pool by finding degenerate oligos in the pool. As previously shown, OligoCompressor reduced a recently published NSR pool of 408 oligos to 262 degenerate oligos[13]. Second, we can avoid brute-force selection of NSR primers and instead design pools of hexamers with optimal coverage and uniformity. The oligos in brute-force pools are scored individually and may not represent the best overall pool when combined. Current NSR pools are designed only to maximize the number of binding sites in the transcript, leading to skewed coverage of transcripts. We developed a multifaceted reward function that scores NSR primer pools using five criteria:

1. *Specificity*. Each NSR primer is compared to hexamers in rRNA and tRNA genes, sequencing adapters, and the other NSR primers. Any NSR candidate that contains these sequences receives a reward of zero.
2. *Gene count*. The agent receives a reward for any gene hit at least once by an oligo in the pool.
3. *Total hits*. The agent is rewarded for maximizing the total number of hits across the transcriptome.
4. *Intergene uniformity*. The agent is rewarded for placing the same number of hits on each gene.
5. *Intragene uniformity*. The agent is rewarded for uniformly distributing hits across the length of each gene.

Inter- and intragene uniformity are quantified by the distribution uniformity, lower quartile (DULQ) score (Figure 3A)[18]:

**Figure 3:**
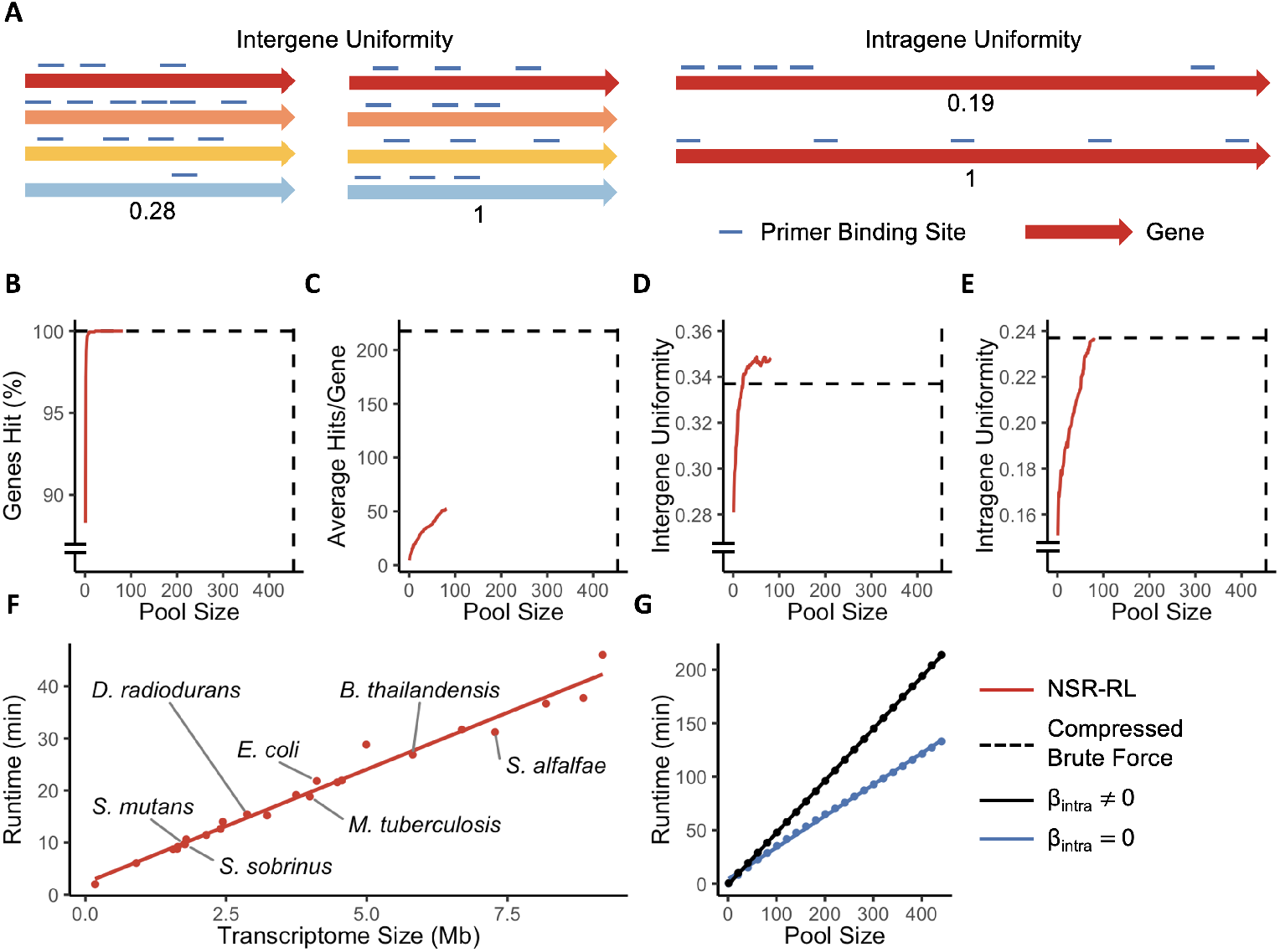
NSR-RL creates hexamer pools using a multivariate reward function. **A**. Intergene uniformity measures the distribution of the hits per gene. Intragene uniformity measures the distribution of hits across the length of each gene. Both uniformity scores range from [0, 1]. NSR hexamer libraries produced by NSR-RL were compared to a pool of 453 hexamers produced by a standard brute-force approach and compressed by OligoCompressor. The libraries were compared across four criteria: the number of unique genes hit at least once (**B**), the total number of hits (**C**), intergene uniformity (**D**), and intragene uniformity (**E**). The dashed black lines show the performance of the brute-force pool, and the solid red lines show the performance of the NSR-RL pool as each hexamer is added to the pool. NSR-RL hit every target with increased intergene uniformity and equivalent intragene uniformity with only 100 oligos. **F**. NSR-RL’s runtime was measured for pools designed to target bacteria with transcriptomes between 0.17 Mb and 9.2 Mb in size. **G**. Quantifying intragene uniformity requires calculating the gaps between all hits on each transcript. Consequently, the runtime of NSR-RL decreases when intragene uniformity is removed from the reward function by setting the associated weight *β*_intra_ = 0.

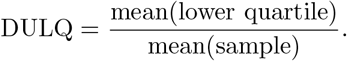

The DULQ is bounded between zero (all hits at a single location) and one (perfect uniformity). Only genes with at least one hit are used to calculate the DULQ. The total reward is the weighted sum of the individual criteria:

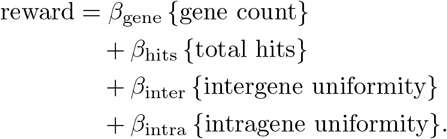

Users can change the weights to emphasize certain criteria when designing primer pools.

We tested the performance of our RL-guided NSR primer design program, called NSR-RL, by designing primer pools using varying weights in the reward function. We compared the NSR-RL pools to primer pools designed using a standard brute-force approach. Both pools targeted the 1.76 Mb *Streptococcus mutans* transcriptome. Changing the reward function weights prioritizes different design criteria. For example, if we are only interested in designing a pool that hits every gene at least once, we can do so by zeroing out the other terms in the reward function. NSR-RL can design a pool that hits every gene using only 10 oligos. A brute-force approach requires 453 oligos to hit every gene even after the final pool is compressed using OligoCompressor.

Rather than minimize the number of oligos, we can use NSR-RL to design a fixed-size pool with improved coverage or uniformity. We used NSR-RL to design a pool containing 100 oligos with nonzero weights for all four criteria in the reward function. The resulting pool exceeded the performance of the compressed brute-force pool (Figure 3B-E). The NSR-RL pool hit every gene in the *S. mutans* transcriptome after only 22 oligos. The NSR-RL pool also placed an average of 993 hits per oligo while the brute-force pool placed an average of 910 hits per oligo. Note that it is impossible to generate more total hits than the brute-force designed pool since the brute-force pool includes all hexamers that are not found in the rRNA or other “unallowable” genes. While the NSR-RL pools contain fewer total hits, the hits are distributed more evenly across the transcriptome as measured by intergene uniformity. Interestingly, we observed that the intergene uniformity score quickly approached a maximum but then oscillated near this value as new oligos were added to the pool. The oscillations indicate that NSR-RL added new oligos that improved the scores of other terms at the expense of intergene uniformity, and vice-versa. The NSR-RL pool’s intragene uniformity matched the performance of the brute-force pool. Users can tune the reward function’s weights to produce NSR primer pools that prioritize either the number of genes hit, total hits, or uniformity. In addition, users can easily add terms to the reward function or create a custom reward to design specialized pools.

NSR-RL’s runtime increases linearly with the size of the problem. We generated NSR pools containing 30 hexamers for an assortment of bacterial transcriptomes ranging between 0.17–9.2 Mb in size. We observed that the algorithm’s runtime scaled linearly with each species’ transcriptome size (Figure 3F). The NSR-RL runtime also increases linearly with the number of oligos in the final pool. The amount of computation required depends heavily on the structure of the reward function. In particular, calculating the intragene uniformity score requires measuring the hit positions of every simulated oligo and calculating the gap distances between each hit position along the length of every gene. Pools designed with reward functions that include intragene uniformity took approximately 50% longer to generate (Figure 3G). We implemented a bypass to skip these calculations if the user is not interested in intragene uniformity, i.e. when the user sets *β*_intra_ = 0.

## 3 Discussion

OligoRL is a general framework designing optimal pools of oligonucleotides. It requires only a reward function and constraints on the action space for each problem; both can be implemented as “black-box” functions. This study used the OligoRL framework to solve three distinct problems. First, CutFreeRL designs pools of DNA barcodes free of restriction sites. CutFreeRL finds near-optimal pools and scales linearly with the problem size, while the original Cut-Free algorithm has exponential scaling. The second tool, OligoCompressor, collapses large, non-degenerate oligo pools into smaller degenerate ones. OligoCompressor works best with large pools of short sequences such as hexamers used for RNA-seq library generation. The smaller pools are less expensive to synthesize, allowing researchers to incorporate base modifications, such as 5’ biotinylation, to their pools. Finally, the NSR-RL tool finds hexamer pools using a complex, multivariate reward function. The resulting pools maximize the number and quality of hits across an entire transcriptome while avoiding unwanted transcripts from rRNA and tRNA genes. Pools designed using NSR-RL require fewer oligos and have better intergene uniformity than pools designed using published brute-force design strategies[4].

The rollout algorithms used in OligoRL have been applied to numerous combinatorial optimization problems. Rollout is a stochastic optimization technique that is not guaranteed to return an optimal solution after a single iteration. One option is to use rollout results to incrementally improve the policy; however, this approach is computationally expensive and the improvements after each iteration can be small. Our results indicate that OligoRL returns near-optimal solutions with a single pass (Figures 1E and 2E). Another option is to exploit the stochasticity of rollout and re-run OligoRL several times with different random number seedings. Stochastic optimizers often include multiple restarts and return the best solution. Indeed, re-running the algorithms in this study produced solutions that were better or worse than our reported results.

All reinforcement algorithms require tuning for optimal performance. For NSR-RL, changing the relative weights of the reward function can impact both solution quality and runtime. Users will also need to balance the time spent finding a single solution (by changing the number of rollout simulations) with re-running the algorithm with different random number seeds. Increasing the number of simulations will improve the agent’s confidence in the expected reward estimates. We found that 100 simulations under each action provided the best results without excessive runtime. Fortunately, the runtime of OligoRL algorithms scales linearly with the number of simulations, so users can easily estimate runtimes based on previous results.

OligoRL uses true “black-box” reward functions. The quality of a candidate oligo pool can be measured using simple algebraic expressions (like degeneracy of the pool) or complex calculations performed by external software packages (such as genome-wide sequence aligners). NSR-RL has a complex, multifactorial reward function, and calculating rewards makes up the majority of the algorithm’s runtime. Researchers with computationally intensive reward functions may consider approximating the reward with a simpler function. Performing more rollout simulations with a less accurate reward may yield better solutions than fewer simulations with better reward estimates.

OligoRL works best when finding optimal solutions from a large set of valid solutions. When the pool of valid solutions shrinks, the nature of the design problem shifts from finding optimal solutions to finding valid solutions that satisfy the problem’s constraints. Rollout, and therefore OligoRL, performs better at optimization than constraint satisfaction. When valid solutions are difficult to find, OligoRL explores many dead-end solutions with poor rewards. For example, instructing NSR-RL maximize total hits leads to states where there are only a few valid hexamers left. In this scenario, OligoRL randomly samples many hexamers but often fails to find the few valid ones. The invalid simulations do not provide useful information to the agent since all invalid actions appear equally poor. Conversely, when nearly all solutions are valid, OligoRL quickly determines good actions for each state since every simulation provides information about an action.

OligoCompressor and NSR-RL find sets of oligos with differing degeneracy. Some wet-lab protocols suggest oligo pools with equimolar concentrations, so experimenters should be careful to mix the oligos in proportion to their degeneracy. The added mixing complexity is a trade-off for the savings gained when using these tools.

Artificial intelligence (AI) is increasingly used in genomic data analysis. This study highlights how AI can improve data collection by optimizing reagents for complex genomic assays. Multiplexed genomic screens need to target thousands of genes while adhering to a large set of biochemical constraints. Such design problems explode combinatorially and are a daunting challenge for traditional optimization techniques. OligoRL demonstrates how reinforcement learning algorithms can simplify complex design problems and find computationally tractable solutions to black box problems. We hope this work will increase the use of AI for designing complex experiments.

## 4 Methods

### 4.1 Implementation

OligoRL and all simulation codes are available as a Julia package at http://jensenlab.net/tools. Simulations were run using Julia version 1.2.8 on a 16-core 3.2 GHz AMD Threadripper processor with 48 Gb of RAM (for OligoCompressor and NSR-RL) or a dual-core 1.6 GHz Intel i5 processor with 8 Gb of RAM (for CutFree and CutFreeRL). The original CutFree algorithm was executed in R version 3.6.2 using Gurobi 9.1.1.

The rollout algorithm used in OligoRL can be parallelized at either the action or simulation level. For example, when simulating the reward for a single base, each simulation can be executed in parallel by a separate thread. This study used Julia’s multithreading tools to perform parallel computations on a multicore processor. The code can also be configured for a cluster computing environment where parallel simulations execute on separate machines.

### 4.2 CutFreeRL Algorithm

CutFreeRL generates a single degenerate oligo that is free of user-specified restriction sites. The specified restriction sites can vary in length up to the length of the final oligo. If any restriction sites are non-palindromic, the reverse complement is added to the list of restriction sites. The user defines the length *L* of the oligo as well as the acceptable base codes that can be used at each position. (Some oligo synthesis companies restrict the use of certain codes.)

An empty oligo is initialized with *L* unspecified positions. Starting at the first position (1), a single candidate base code *a* is selected from the set *A*(*s*_1_) of allowed codes at position 1. The algorithm simulates ahead to form a candidate oligo that begins with code *a* at position 1 followed by randomly selected codes through the end of the oligo (positions 2 through *L*). At each position *i*, the code is selected from the set *A*(*s*_*i*_) of allowed codes at that position. As codes are selected, the algorithm checks that the newest code does not create a restriction site in the candidate oligo. A (backwards) sliding horizon is used to checking the oligo for restriction sites, with the horizon equal to the length of the longest restriction site. Any codes beyond the window have already been checked for restriction sites when those codes were selected. If all of the allowed codes create a restriction site, the simulation stops and oligo is terminated early. At the end of each simulation (i.e. after a single candidate oligo has been built), the reward function returns the log_2_ degeneracy of the candidate. Any prematurely terminated oligo candidate is assigned a log_2_ degeneracy of −100, a prohibitively large penalty.

CutFreeRL performs many simulations (100–1000) for each allowed base code *a* at the first position 1. The overall reward for code *a* is the average reward of the individual simulations. The code with the highest average reward is selected for position 1, and the algorithm moved to the next position 2. The entire simulation procedure is repeated starting at position 2, and so on until codes have been selected for all *L* positions.

Benchmarking experiments were performed on groups of restriction sites selected randomly from the set of all commercially available, palindromic, hexameric restriction enzymes in REBASE[12]. The groups of restriction sites were blocked from 20 bp oligos using both CutFreeRL and the original CutFree algorithm. Randomly selected sets of five restriction sites were used for experiments that varied the oligo length. All base codes were allowed at every position, and CutFreeRL performed 1000 rollout simulations per action. The following linear model was used to compare the decrease in degeneracy as additional restriction sites are blocked:

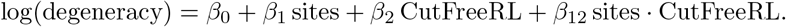

A *t*-test on the coefficient *β*_12_ assessed the significance of the interaction between the number of sites and the choice of algorithm.

### 4.3 OligoCompressor Algorithm

OligoCompressor reduces a large set of non-degenerate oligos (the target pool) into a smaller set of degenerate ones (the compressed pool). OligoCompressor applied a semi-greedy approach that begins by designing a degenerate oligo that matches the largest number of oligos in the target pool. The resulting oligo is added to the compressed pool and the matched sequences are removed from the target pool. OligoCompressor repeats this process until all oligos in the target pool have been matched by an oligo in the compressed pool.

OligoCompressor begins with an empty oligo equal to the length of the oligos in the target pool. Rollout is used to select degenerate bases for each position given the expected future rewards. If at any point a candidate oligo contains a sequence that is not in the original target pool, the simulation is stopped prematurely, and the candidate receives a score of zero; otherwise, the candidate’s score is the number of matched oligos in the target pool. To check if a candidate oligo contains a non-target sequence, the algorithm checks that the degeneracy of the candidate does not exceed the number of matches in the original target pool.

As mentioned previously, OligoCompressor updates the action space of allowed bases as it builds candidate oligos (Figure 2B). For example, if the algorithm selects code `T` at position 1, the action spaces for the remaining positions are limited to only those codes that appear in oligos that begin with `T`. If code `A` is selected for position 2, the action space for positions 3 and beyond are further resticted to bases that appear in target oligos beginning with `TA`. Shrinking the action spaces prevents unnecessary simulations by skipping codes that are guaranteed to end prematurely when the candidate does not match any target oligos. It is especially useful near the end of the algorithm when the few remaining target sequences need to be added individually to the compressed pool.

To benchmark OligoCompressor (Figure 2C,D), pools of random hexamers of size 50–4,096 were sampled without replacement from the set of all possible hexamers. To test compression efficiency (Figure 2e), pools of 10 degenerate oligos were constructed that contained 30 to 150 unique oligos. Each degenerate oligo was generated by sampling bases as Poisson random variable ranging from degeneracy 1 (`A`, `T`, `G`, or `C`) to degeneracy 4 (`N`). The degeneracy of each pool was tuned by changing the Poisson rate parameter (*λ*). Each pool of 10 degenerate oligos was expanded and then re-compressed using OligoCompressor. All benchmarking experiments used 100 rollout simulations per action; no differences in compression efficiency were observed using 1000 simulations per action.

### 4.4 NSR-RL Algorithm

NSR-RL designs Not-So-Random primer pools for RNA-seq library preparation and other multiplex genomic assays. Users supply two sequence files containing 1.) “targeted” transcripts that should be targeted by the NSR primers, and 2.) “unallowed” transcripts to avoid, e.g. transcripts from rRNA and tRNA genes. The user also specifies the number of NSR primers to create and the length of the primers (the default is hexamers).

NSR-RL builds oligos using rollout with dynamic action spaces as described for the OligoCompressor. Candidate oligos are assigned a reward of zero if they hit any unallowed transcript. Palindromic candidates are also assigned a reward of zero since palindromic reverse transcription primers may self-anneal during amplification. Non-palindromic candidate oligos that miss the unallowed transcripts are scored by the multifaceted reward function. The first three terms in the reward are calculated by counting the number of times the oligo hits each targeted sequence. First, the gene count term is the number of genes that are hit at least once. Second, the total hits term is sum of all hits across the transcriptome. Third, the intergene uniformity score is calculated using the DULQ score of all of the hit counts. Calculating the fourth term in the reward, intragene uniformity, requires the gaps between hits to calculate the DULQ for each transcript. Transcripts with a more uniform gap distance distribution will score higher than transcripts with different sized gaps. The overall intragene uniformity score is the average DULQ across all transcripts. We multiply the inter- and intragene uniformity scores by the number of targets, *n*_targets_, to place these rewards on a similar scale as the other terms. The target count and uniformity terms range from 0 to *n*_targets_, while the total hits is term is unbounded. Each term in the reward function has an associated weight *β*, and the weights can be changed to tune the pools empirically.

After NSR-RL finishes an oligo, the oligo and its reverse complement are added to the list of unallowed sequences to prevent avoid repeats or selecting oligos that could form dimers when the libraries are amplified.

NSR-RL was benchmarked by creating 100 degenerate hexamers targeting the transcriptome of the 1076 Mb transcriptome of *Streptococcus mutans* strain UA159 (Figure 3B–E). Unless otherwise specified, the reward weights were *β*_gene_ = 1, *β*_hits_ = 10^*−*4^, *β*_inter_ = 1, and *β*_intra_ = 1. To compare NSR-RL runtime with transcriptome size (Figure 3F), 30 degenerate hexamers were designed to target the transcriptomes of 25 species of bacteria (Supplementary File 1).

## 5 Acknowledgements

This work was supported by the National Institutes of Health (grant GM138210) and the Laboratory Directed Research and Development (LDRD) Program of Sandia National Laboratories (contract 2093481) to PAJ. Sandia National Laboratories is a multi-mission laboratory managed and operated by National Technology & Engineering Solutions of Sandia, LLC, a wholly owned subsidiary of Honeywell International Inc., for the U.S. Department of Energy’s National Nuclear Security Administration under contract DE-NA0003525. BMD is supported in part by an Illinois Distinguished Fellowship.

## References

[1] M. Hendling, S. Pabinger, K. Peters, N. Wolff, R. Conzemius, and I. Barišić, “Oli2go: an automated multiplex oligonucleotide design tool,” Nucleic Acids Research, vol. 46, no. Web Server issue, W252, Jul. 2018. doi: 10.1093/NAR/GKY319. [Online]. Available: /pmc/articles/PMC6030895/%20/pmc/articles/PMC6030895/?report=abstract%20 https://www.ncbi.nlm.nih.gov/pmc/articles/PMC6030895/.

[2] M. Hendling and I. Barišić, “Insilico Design of DNA Oligonucleotides: Challenges and Approaches,” Computational and Structural Biotechnology Journal, vol. 17, pp. 1056–1065, Jan. 2019, ISSN: 2001-0370. doi: 10.1016/J.CSBJ.2019.07.008.

[3] A. J. Storm and P. A. Jensen, “Designing Randomized DNA Sequences Free of Restriction Enzyme Recognition Sites,” Biotechnology Journal, vol. 13, no. 1, Jan. 2018, ISSN: 18607314. doi: 10.1002/biot.201700326. [Online]. Available: https://pubmed.ncbi.nlm.nih.gov/28865135/.

[4] C. D. Armour, J. C. Castle, R. Chen, T. Babak, P. Loerch, S. Jackson, J. K. Shah, J. Dey, C. A. Rohl, J. M. Johnson, and C. K. Raymond, “Digital transcriptome profiling using selective hexamer priming for cDNA synthesis,” Nature Methods 2009 6:9, vol. 6, no. 9, pp. 647–649, Aug. 2009, ISSN: 1548-7105. doi: 10.1038/nmeth.1360. [Online]. Available: https://www.nature.com/articles/nmeth.1360.

[5] M. Vignali, C. D. Armour, J. Chen, R. Morrison, J. C. Castle, M. C. Biery, H. Bouzek, W. Moon, T. Babak, M. Fried, C. K. Raymond, and P. E. Duffy, “NSR-seq transcriptional profiling enables identification of a gene signature of Plasmodium falciparum parasites infecting children,” The Journal of Clinical Investigation, vol. 121, no. 3, p. 1119, Mar. 2011. doi: 10.1172/JCI43457. [Online]. Available: /pmc/articles/PMC3046638/%20/pmc/articles/PMC3046638/?report=abstract%20 https://www.ncbi.nlm.nih.gov/pmc/articles/PMC3046638/.

[6] O. Arnaud, S. Kato, S. Poulain, and C. Plessy, “Targeted reduction of highly abundant transcripts using pseudo-random primers,” BioTechniques, vol. 60, no. 4, pp. 169–174, Apr. 2016. doi: 10.2144/000114400. [Online]. Available: http://www.BioTechniques.com.

[7] A. Cornish-Bowden, “Nomenclature for incompletely specified bases in nucleic acid sequences: recommendations 1984.,” Nucleic Acids Research, vol. 13, no. 9, p. 3021, May 1985. doi: 10.1093/NAR/13.9.3021. [Online]. Available: https://www.ncbi.nlm.nih.gov/pmc/articles/PMC341218/.

[8] A. Untergasser, I. Cutcutache, T. Koressaar, J. Ye, B. C. Faircloth, M. Remm, and S. G. Rozen, “Primer3—new capabilities and interfaces,” Nucleic Acids Research, vol. 40, no. 15, e115, Aug. 2012. doi: 10.1093/NAR/GKS596. [Online]. Available: /pmc/articles/PMC3424584/%20/pmc/articles/PMC3424584/?report=abstract%20https://www.ncbi.nlm.nih.gov/pmc/articles/PMC3424584/.

[9] D. P. Bertsekas, Reinforcement Learning and Optimal Control, 1st Ed. Athena Scientific, 2019, ISBN: 978-1-886529-39-7.

[10] D. P. Bertsekas, Rollout, Policy iteration, and Distributed Reinforcement Learning. Athena Scientific, 2020, ISBN: 978-1-886529-07-6.

[11] R. Bellman, “A Markovian Decision Process,” Journal of Mathematics and Mechanics, vol. 6, no. 5, pp. 679–684, 1957.

[12] R. J. Roberts, T. Vincze, J. Posfai, and D. Macelis, “REBASE—a database for DNA restriction and modification: enzymes, genes and genomes,” Nucleic Acids Research, vol. 43, no. D1, pp. D298–D299, Jan. 2015, ISSN: 0305-1048. doi: 10.1093/NAR/GKU1046. [Online]. Available: https://academic.oup.com/nar/article/43/D1/D298/2436339.

[13] T. Hayashi, H. Ozaki, Y. Sasagawa, M. Umeda, H. Danno, and I. Nikaido, “Single-cell full-length total RNA sequencing uncovers dynamics of recursive splicing and enhancer RNAs,” Nature Communications 2018 9:1, vol. 9, no. 1, pp. 1–16, Feb. 2018, ISSN: 2041-1723. doi: 10.1038/s41467-018-02866-0. [Online]. Available: https://www.nature.com/articles/s41467-018-02866-0.

[14] A. J. Westermann, S. A. Gorski, and J. Vogel, “Dual RNA-seq of pathogen and host,” Nature Reviews Microbiology 2012 10:9, vol. 10, no. 9, pp. 618–630, Aug. 2012, ISSN: 1740-1534. doi: 10.1038/nrmicro2852. [Online]. Available: https://www.nature.com/articles/nrmicro2852.

[15] R. Sooknanan, J. Hitchen, A. Radek, and J. Pease, “Superior rRNA Removal for RNA-Seq Library Preparation,” Journal of Biomolecular Techniques : JBT, vol. 23, no. Suppl, S57, 2012. [Online]. Available: /pmc/ articles/PMC3630600/?report=abstract%20https://www.ncbi.nlm.nih.gov/pmc/articles/PMC3630600/.

[16] F. J. Stewart, E. A. Ottesen, and E. F. DeLong, “Development and quantitative analyses of a universal rRNA-subtraction protocol for microbial metatranscriptomics,” The ISME journal, vol. 4, no. 7, pp. 896–907, Jul. 2010, ISSN: 1751-7370. doi: 10.1038/ISMEJ.2010.18. [Online]. Available: https://pubmed.ncbi.nlm.nih.gov/20220791/.

[17] P. H. Culviner, C. K. Guegler, and M. T. Laub, “A simple, cost-effective, and robust method for rrna depletion in rna-sequencing studies,” mBio, vol. 11, no. 2, Mar. 2020. doi: 10.1128/MBIO.00010-20.

[18] C. Burt and S. Styles, Drip and micro irrigation for trees, vines, and row crops: Design and management (with special sections on SDI). 1999, ISBN: 978-0964363427.

